## Supplementary Information for "A fungal metabolic regulator underlies infectious synergism during *Candida albicans*-*Staphylococcus* aureus intra-abdominal co-infection"

### SUPPLEMENTARY METHODS

#### Phenotypic profiling of *C. albicans* strains

Growth was assessed by diluting overnight YPD grown cultures to 1x$10^5$ cells/mL in fresh YPD (or other media as specified), transferring 200 µL to a microtiter plate, and kinetically measuring the OD600 nm at hourly intervals using a BioTek Synergy spectrophotometer. In some experiments, the pH of culture supernatants was assessed by first filter sterilizing the spent medium using a 0.22 µm filter and using a standard laboratory pH meter. Additionally, hyphal growth assays were carried out by diluting (1:100) overnight cultures of YPD grown cells in Roswell Park Memorial Institute 1640 medium (RPMI) and incubating at 37ºC with shaking. Aliquots were removed at 2 and 4 h post-inoculation, transferred to glass slides, and images digitally captured by standard light microscopy. To determine susceptibility to common stressors YPD grown cells were washed 3X and diluted in sterile water. Aliquots containing 5-5x$10^4$ cells were spotted onto YPD plates containing the following stressors: 100 mM CaCl2, 500 mM CaCl2, 100 µg/mL Congo Red, 3% ethanol, 3% glycerol, 10 mM MnCl2, 1.5 M NaCl, 0.05% SDS, 1 M sorbitol, 5 mM H2O2, 1 µg/mL fluconazole, or 5 µg/mL fluconazole. Susceptibility to pH was similarly tested in YPD medium adjusted to pH 3.5 or 8.5. Plates were incubated at 30ºC and growth assessed at 24 h. Temperature sensitivity was determined by plating onto YPD and incubating at 37ºC or 42ºC for 24 h. Images were acquired digitally using a Gel Doc XR Imaging System.

#### α-toxin ELISA protocol

Microtiter plates were coated with 50 µL 0.1 µg/mL anti-α-toxin antibody MEDI4893* diluted in ELISA/ELISPOT coating buffer (Invitrogen). Plates were incubated at 4°C overnight, washed with PBS-Tween 20 (PBS-T), and blocked with SuperBlock (Pierce) for 1 h at room temperature. Then,

50 µL filter-sterilized culture supernatant, peritoneal lavage fluid, or clarified organ homogenate were serially diluted in PBS and plated along with native α-toxin as the standard. Plates were incubated for 1 h at room temperature and then washed with PBS-T. Affinity-purified rabbit polyclonal anti-α-toxin antibody (2 µg/mL) was added for 1 h at room temperature. Plates were washed with PBS-T. AffiniPure horseradish peroxidase (HRP)-coupled goat anti-rabbit IgG detection antibody (Jackson ImmunoResearch) was diluted 1:10000 and incubated for 1 h at room temperature. After extensive washing with PBS-T, 100 µL 3,3',5,5'-Tetramethylbenzidine (TMB) substrate was added, and plates were incubated in the dark for 10 min. 100 µL of 0.2 N $H_2SO_4$ was immediately added to stop the reaction. Plates were read at 450 nm using a BioTek Synergy H1. Experimental concentrations were extrapolated to the standard curve. Culture supernatants or in vivo samples from the *S. aureus* Δ*hla* strain were blank subtracted to account for any nonspecific antibody binding to shed protein A.

**Site-directed mutagenesis**

Construction of the oligomerization-deficient H35L α-toxin mutant was performed using Platinum SuperFi I (ThermoFisher) following the manufacturer's site-directed mutagenesis protocol. Primers H35L-Superfi-F and H35L-Superfi-R **(Supplementary Table 3)** were used to PCR amplify previously constructed pSK-hla [1]. Amplified DNA was digested with DpnI to remove residual template DNA and transformed into the *dcm*-deficient *E. coli* strain IM08B to yield plasmid pSK-H35L (**Supplementary Table 4**). pSK-H35L was electroporated into *S. aureus* NE1345 Δ*hla* and selected on TSA with 10 µg/mL chloramphenicol to create strain Δ*hla*-p*hla*$^{H35L}$. Sanger sequencing using primers hlaSeqF and hlaSeqR was performed to verify plasmid integrity.

**Construction of the *agr* P3 luciferase reporter**

To construct a P3-luciferase reporter plasmid, the *S. aureus agr* P3 promoter was PCR amplified from synthetic DNA (Integrated DNA Technologies) using primers SynthGeneF and SynthGeneRv4 **(see Supplemental Table 5).** This PCR product and plasmid pMV306G13+Lux

(Addgene, #26160) containing the *luxABCDE* operon were digested with NotI and NcoI, ligated, and transformed into *E.coli* DH5-α to yield pOLux1 using the method previously described [2]. The entire P3-*luxABCDE* fragment from pOLux1 was PCR amplified using primers luxABCDE-Fv2-EcoRI and luxABCDE-R-NarI. This product, along with plasmid pMK4, were digested with EcoRI and NarI. Digests were ligated and transformed into *E. coli* DH5-α, yielding plasmid pOLux2. The constitutive *S. aureus* promoter *sarA* P1 was synthesized (IDT) and amplified with primers SynthGeneF and SynthGeneRv4 (**Supplementary Table 5**). This product and pOLux2 were digested with Acc65I, ligated, and transformed into *E. coli* DH5-α, yielding pOLux3 (**Supplementary Table 4**). Correct orientation of the *sarA* P1 promoter in pOLux3 was confirmed by PCR amplification with primers sarAP1-FAcc65I and luxC-DET-R. pOLux3 was transformed into the *dcm*-deficient *E. coli* strain IM08B, as previously described [2]. After confirmatory digestion and sequence integrity verified, plasmids were transformed into *S. aureus* JE2 competent cells and plated on brain heart infusion agar containing 10 μg/mL chloramphenicol to yield strain *S. aureus*(pOLux).

**_ZCF13_ disruption plasmid construction**

Plasmids for *ZCF13* gene disruption were constructed using plasmid pBSS2, containing the *SAT1*-flipper disruption cassette [3]. The 5' flanking region of the *ZCF13* ORF was amplified from SC5314 genomic DNA using primers ZCF13-FF-KpnI and ZCF13FR-ApaI, digested with KpnI and ApaI, and ligated into pBSS2 at the KpnI and ApaI restriction sites to create pBSS-2ZCF13-F. The 3' flanking region of the *ZCF13* ORF was amplified from SC5314 gDNA using primers ZCF13-RF-NotI and ZCF13-RR-SacI, digested with NotI and SacI, and ligated into pBSS2-ZCF13-F at the NotI and SacI restriction sites to create pBSS2-ZCF13-FR (**Supplementary Table 4**).

**_ZCF13_ mutant construction**

Plasmid pBSS2-ZCF13-FR was digested with KpnI and SacI and the linear fragment was transformed into SC5314 using a standard lithium acetate protocol, with some modifications [4, 5]. An overnight culture of SC5314 was diluted 1:200 in 5 mL YPD and grown for 6 h at 30°C with shaking at 200 rpm and washed by centrifugation using sterile water. Cells were then resuspended in 1X TELiAc buffer (100 mM LiAc, 1X TE) and 1 μg of linearized pBSS2-ZCF13-FR was added to 50 μL cell suspension. Single-stranded salmon sperm carrier DNA (5 μL) was added along with 300 μL 40% PEG. The transformation mix was incubated at 30°C for 30 min with agitation every 10 min. Cells were heat-shocked at 42°C for 15 min and then allowed to recover in YPD at 30°C with shaking at 200 rpm for 4-6 hours. Cells were pelleted and resuspended in 1X TE and plated on YPD+200 μg/mL nourseothricin (YPD+200NAT). Plates were incubated for up to 2 days at 30°C. Colonies were grown overnight in 1 mL YPM (YP+2% maltose) to induce cassette excision. Cells from YPM cultures were washed with PBS and then diluted to $10^3$ cells in 1X TE buffer. 100 μL cells were plated onto YPD+25 μg/mL nourseothricin (YPD+25NAT) and incubated at 30°C for 24 hours. Small and medium-sized colonies were patched onto YPD and YPD+200NAT and incubated at 30°C for 24 hours. Colonies that grew on YPD but not YPD+200NAT were confirmed by PCR to have *ZCF13* disrupted using primers ZCF13-FLPINTF + ZCF13-DET-R and ZCF13-FLPINTR + ZCF13-DET-F. The transformation procedure was repeated to disrupt the second *ZCF13* allele. Homyzogous deletion of *ZCF13* was verified with primers ZCF13-DET-F + ZCF13-DET-R.

**_ZCF13_ revertant construction**

One copy of *ZCF13* was inserted into the neutral locus *NEUT5L* of *zcf13*Δ/Δ using the shuttle vector pDUP3. The *ZCF13* 5' UTR, ORF, and 3' UTR was PCR amplified from SC5314 using primers ZCF13-AMPF-SmaI and ZCF13-AMPR-NotI. The PCR product and plasmid pDUP3 were digested with SmaI and NotI and ligated to create pDUP3-ZCF13 (**Supplementary Table 4**). After transformation into DH5α and plasmid recovery, pDUP3-ZCF13 was linearized by digesting with SfiI and then transformed into *zcf13*Δ/Δ following the lithium acetate protocol described above.

Revertant colonies were selected for by growth on YPD+200NAT and confirmed by PCR using primers NAT1INTF + NEUT5LAMPF and NEUT5LAMPR + ZCF13-DET-R.

**Construction of *rbk1*Δ/Δ**

To construct the *rbk1*Δ/Δ strain, linearized *SAT1*-flipper and *CaHygB*-flipper plasmids (**Supplementary Table 4**) were amplified with RBK1CC9KO-F and RBK1CC9KO-R primers to make the disruption repair templates harboring nourseothricin or hygromycin resistance cassettes, respectively as described [6, 7]. Before the transformation, *C. albicans* competent cells were treated with 1X Tris-EDTA and 0.1 M lithium acetate (pH 7.5) for 1 h at 37°C followed by addition of 25 mM dithiothreitol for another 30 minutes. Cells were then washed with sterile water and 1 M sorbitol and resuspended in residual sorbitol. The cells were mixed with premade ribonucleoprotein (RNP) complex containing universal tracrRNA, Cas9 protein (2 μg), *RBK1* gene-specific guide RNAs (crRBK1up and RBK1down), and 1 μg of each PCR generated repair template. The transformation was performed by a single pulse at 1.8 kV using the Gene Pulser Xcell Electroporation System (Bio-Rad). Immediately, cells were incubated at 30°C for 4-6 hours in YPD medium and plated on YPD+200NAT and 600μg/mL hygromycin B (YPD+600HygB). Selected colonies were transferred to a yeast-peptone medium with 2% maltose (YPM) for overnight culture to excise the cassettes. Cells were then selected in YPD agar media with nourseothricin (25 μg/mL) and hygromycin (75μg/mL). Genetic integration and excision of the resistance cassettes were confirmed phenotypically by growth on YPD, YPD+200NAT, and YPD+600HygB solid media. Correct genomic integration and loss of *RBK1* was confirmed by PCR amplification of genomic DNA using primers RBK1INTF+FLPINTR, RBK1INTR+FLPINTF, and RBK1DETF+RBK1DETR **(Supplementary Table 3)** [6, 7].

**Construction of the *rbk1*Δ/Δ+*RBK1* strain**

To make the *rbk1*Δ/Δ+*RBK1* strain, part of the *NEUT5L* homology region, *RBK1* promoter and ORF, and ADH1terminator were PCR amplified from either SC5314 gDNA or pDIS3-tAHD1 plasmid using primer pairs NEUT5homology-pDISF+Nt5ADHup-UNIVOL-R, RBK1_Pr_OL-F+RBK1_ORF-OL-R, and ADH1t-UNIVOL-F+ NEUT5homology-pDISR (**Supplementary Table 3**) as described previously[7]. Overlap extension PCR of the amplified products was performed and a final amplification of the sewn product was achieved using NEUT5homology-pDISF and NEUT5homology-pDISR primers to make the repair template. The transformation was performed using the previously mentioned CRISPR-Cas9 protocol, except the crNEUT5L-up_pDIS3 crRNA was used during the preparation of RNP assembly. Transformants were selected on YPD-200NAT. Correct genomic integration and gene reversion were confirmed by using primers NAT1INTF+NEUT5LAMPR and RBK1DETF+RBK1DETR **(Supplementary Table 3)** [6, 7].

**Construction of *RBK1*, *HGT7*, or double overexpression strains**

To construct overexpression strains *zcf13*Δ/Δ+Pr*TEF1-RBK1* or *zcf13*Δ/Δ+Pr*TEF1-HGT7,* the coding sequence of *RBK1* and *HGT7* was PCR amplified from SC5314 genomic DNA using primer pairs RBK1_ORF-F-SalI+ RBK1_ORF-R-MluI or HGT7_ORF-F-SalI+ HGT7_ORF-R-MluI, respectively. Plasmid pKE4 and PCR amplified products were then digested with restriction enzymes SalI+MluI and ligated to make pKE4-*RBK1* or -*HGT7* (**Supplementary Table 4**)*.* Part of *the* NEUT5 homology region and ADH1terminator were PCR amplified from pDIS3-tAHD1 as previously described. Plasmids pKE4-Pr*TEF1-RBK1* or -*HGT7* were amplified using primers PrTEF1OLF and tADH1OLR. The overlap extensions of the amplified products and subsequent steps were performed exactly as described above. Genotypes were confirmed by using primer pairs NAT1INTF+NEUT5LAMPR and either RBK1DETF+NAT1_INTR or HGT7DETF+NAT1_INTR.

The *RBK1* and *HGT7* double overexpression strain (*zcf13*Δ/Δ+Pr*TEF1-RBK1-HGT7*) was constructed using strain *zcf13*Δ/Δ+Pr*TEF1-HGT7.* Part of the *NEUT5* homology region, Pr*TEF1-RBK1*, and the hygromycin resistance cassette were amplified from either pDIS3-tAHD1, pKE4-Pr*TEF1-RBK1,* or pHygR-FLP by using primer pairs NEUT5homology-pDISF+Nt5ADHup-UNIVOL-R, PrTEF1OLF+ADHterm-OL-R, PrHygR-OL-F+tHygR-OL-R, or NEUT5L_OL-F+

NEUT5homology-pDISR, respectively as described above. Transformants were selected on YPD-200NAT or YPD-600HygB medium. The genotype was confirmed using primers RBK1DETF+ADH13SEQR and NEUT5LAMPF+ CaHygB-SEQR.

**RNA-Seq library preparation**

For *C. albicans*, 1 μg RNA per sample was used for library preparation. Sequencing libraries were generated using NEBNext®UltraTMRNA Library Prep Kit for Illumina® following manufacturer's recommendations. Briefly, mRNA was purified from total RNA using poly-T oligo-attached magnetic beads. Fragmentation was carried out using divalent cations under elevated temperature in NEBNext First Strand Synthesis Reaction Buffer. First strand cDNA was synthesized using random hexamer primers and M- MuLV Reverse Transcriptase (RNase H-). Second strand cDNA synthesis was subsequently performed using DNA Polymerase I and RNase H. Remaining overhangs were converted into blunt ends via exonuclease/polymerase activities. After adenylation of 3' ends of DNA fragments, NEBNext Adaptor with hairpin loop structures were ligated to prepare for hybridization. Size-selected cDNA fragments of preferentially 150~200 bp in length were purified with AMPure XP system. Then 3 μl USER™ enzyme (NEB, USA) was used with size-selected, adaptor-ligated cDNA and PCR was performed with Phusion High-Fidelity DNA polymerase. Lastly, PCR products were purified (AMPure XP system) and library quality was assessed on the Agilent Bioanalyzer 2100 system.

For *S. aureus,* ribosomal RNA was removed from total RNA using Illumina Ribo-Zero Plus rRNA depletion Kit according to the manufacturer's instructions, followed by ethanol precipitation. During the second strand cDNA synthesis, dUTPs were replaced with dTTPs in the reaction buffer. The directional library was ready after end repair, A-tailing, adapter ligation, size selection, USER enzyme digestion, amplification, and purification. The rest of the sequencing pipeline was similar for both microbes. In brief, samples were clustered by a cBot Cluster Generation System using PE Cluster Kit cBot-HS according to the manufacturer's protocol and were sequenced by generating 150 bp paired-end reads using an Illumina platform.

**RNA-Seq raw data processing**

Raw data in FASTQ format were processed and trimmed by removing adapters, poly-N, and low-quality reads. At the same time, Q20, Q30, and GC content of the cleaned data were calculated and used for downstream analyses. The *C. albicans* SC5314 (C_albicans_SC5314_version_A22-s07-m01-r101_default_genomic.fasta) and *S. aureus* FPR3757 ( 000013465.ASM1346v1.dna.chromosome.Chromosome.fa.gz) reference genomes and gene model annotation files were downloaded from the Candida Genome Database or Ensemble Genomes, respectively. HISAT2 [8] or Bowtie2 [9] were used for *C. albicans* and *S. aureus,* respectively, to build a genome indices and perform alignments (**Supplementary Table 6**).The read numbers mapped to each gene were counted using FeatureCounts [10]. Gene expression levels were estimated by the expected number of Fragments Per Kilobase of transcript sequence per Million base pairs (FPKM) sequenced [11].

**Homology search and alignment of ScGal2 orthologs**

The *S. cerevisiae* Gal2 amino acid sequence was retrieved from the *Saccharomyces* Genome Database and a protein homology search conducted against the *C. albicans* SC5314 genome using NCBI BLAST. Clustal Omega was used to perform a multiple sequence alignment between ScGal2p and the homologous CaHgt7p and CaHgt8p amino acid sequences.

**SUPPLEMENTARY TABLES**

| Strain | Organism | Parent | Relevant Genotype | Reference |
|---|---|---|---|---|
| DH5α | *E. coli* | -- | Amp$^S$; cloning host | (12) |
| IM08B | E. coli | DC10B | Amp$^S$; SA08BΩPN25-hsdS (CC8-1) (SAUSA300_0406) of NRS384 integrated between the essQ and cspB genes | (13) |
| JE2 | *S. aureus* | LAC | Erm$^S$, Cm$^S$ | (14) |
| *S. aureus*(pDB22) | *S. aureus* | JE2 | Erm$^R$, Cm$^R$; contains *agr* P3-GFP reporter plasmid pDB22 | (2) |
| *S. aureus*(pOLux) | *S. aureus* | JE2 | Erm$^R$, Cm$^R$; contains *agr* P3-*luxABCDE* reporter plasmid pOLux3 | this study |
| Δ*hla* | *S. aureus* | JE2 | Erm$^R$; *hla*::*bursa* | (14) |
| Δ*hla*+*phla* | *S. aureus* | JE2 | Erm$^R$, Cm$^R$; contains plasmid pSK-*hla* | (2) |
| Δ*hla*+*phla*$^{H35L}$ | *S. aureus* | Δ*hla* | Erm$^R$, Cm$^R$; contains plasmid pSK-H35L | this study |

**Supplementary Table 1. Bacterial strains constructed or used in this study.** Sensitivity ($^S$) or resistance ($^R$) to the following antibiotics is annotated: ampicillin (Amp), erythromycin (Erm), or chloramphenicol (Cm).

| **Strain name** | **Parent** | **Genotype** | **Reference** |
|---|---|---|---|
| SC5314 | -- | wild-type, reference isolate | (15) |
| TF WT | SN152 | *arg4*Δ/*arg4*Δ *leu2*Δ/*LEU2 his1*Δ/*HIS1 URA3*/*ura3*Δ::imm434 *IRO1*/*iro1*Δ::imm434 | (16) |
| TF *zcf13*Δ/Δ (TF6) | SN152 | *arg4*Δ/*arg4*Δ *zcf13*Δ::*LEU2 zcf13*Δ::*HIS1 URA3*/*ura3*Δ::imm434 *IRO1*/*iro1*Δ::imm434 | (16) |
| *zcf13*-het | SC5314 | *zcf13*Δ:*FRT* / *ZCF13 NEUT5L*/*NEUT5L* | this study |
| *zcf13*Δ/Δ | *zcf13*-het | *zcf13*Δ:*FRT zcf13*Δ:*FRT NEUT5L*/*NEUT5L* | this study |
| *zcf13*Δ/Δ+*ZCF13* | *zcf13*Δ/Δ | *zcf13*Δ:*FRT zcf13*Δ:*FRT neut5l*Δ:Pr*ZCF13*-*ZCF13*-t*ZCF13*/*NEUT5L* | this study |
| *rbk1*Δ/Δ | SC5314 | *rbk1*Δ:*FRT*+ *rbk1*Δ:*FRT*+ *NEUT5L*/*NEUT5L* | this study |
| *rbk1*Δ/Δ+*RBK1* | *rbk1*Δ/Δ | *rbk1*Δ:*FRT*+ *rbk1*Δ:*FRT*+ *neut5l*Δ:Pr*RBK1*-*RBK1*-t*ADH1*/*NEUT5L* | this study |
| *zcf13*Δ/Δ+Pr*TEF1*-*RBK1* | *zcf13*Δ/Δ | *zcf13*Δ:*FRT*+ *zcf13*Δ:*FRT*+ *neut5l*Δ:Pr*TEF1*-*RBK1*-t*ADH1*/*NEUT5L* | this study |
| *zcf13*Δ/Δ-Pr*TEF1*-*HGT7* | *zcf13*Δ/Δ | *zcf13*Δ:FRT+ *zcf13*Δ:*FRT*+ *neut5l*Δ:Pr*TEF1*-*HGT7*-t*ADH1*/*NEUT5L* | this study |
| *zcf13*Δ/Δ-Pr*TEF1*-*HGT7*-*RBK1* | *zcf13*Δ/Δ | *zcf13*Δ:*FRT*+ *zcf13*Δ:*FRT*+ *neut5l*Δ:Pr*TEF1*-*HGT7*-t*ADH1* *neut5l*Δ:Pr*TEF1*-*RBK1*-t*ADH1* | this study |

**Supplementary Table 2.** ***C. albicans*** **strains constructed or used in this study.**

| name | Sequence (5’ → 3’)[a] |
|---|---|
| **Primers** | |
| H35L-Superfi-F | ATGGCATGC***T***CAAAAAAGTATTTTATAG |
| H35L-Superfi-R | TACTTTTTTG***A***GCATGCCATTTTCTTTATC |
| hlaSeqF | GATATGTCTCAACTGCAATATTCTAAATTGACATA |
| hlaSeqR | ACATCATTTCTGAAGTTATCGGC |
| SynthGeneF | CGCAGTTACGGATCAGTCAC |
| SynthGeneRv4 | GGTGCTGCCATGTTCTTTGCT |
| P3lux-F-NotI | TCAGCGGCCGCATTTTAACATAAAAAAATTTACAGTTAAGAATAAAAAACGACTAG |
| P3Lux-R-NcoI | TCACCATGGGATCTCTGTAATCTAGTTATATTAAAACATGCTAAAAGCAT |
| luxABCDE-Fv2-EcoRI | TCAGAATTCATTTTAACATAAAAAAATTTACAGTTAAGAATAAAAAACGACTAG |
| luxABCDE-R-NarI | TCAGGCGCCGATCACCGCGGCCATGAT |
| sarAP1-F-Acc65I | TCAGGTACCCTGATATTTTTGACTAAACCAAATGC |
| luxC-DET-R | GTCACGAATGTATGTCCTGC |
| ZCF13-FF-KpnI | TCAGGTACCAATCAAGCCTCCTGTACCACCACCA |
| ZCF13-FR-ApaI | TCAGGGCCCGCCTGGACTATTTGTCTTATCCATAATCGAT |
| ZCF13-RF-NotI | TCAGCGGCCGCGTGATGAAATTTTGGATCCTGTTGCTTG |
| ZCF13-RR-SacI | TCAGAGCTCGGCATGTTGTTGCTTTAGTGTCAGG |
| ZCF13-AMPF-SmaI | TCACCCGGGGACATCTCTCATTTGGTATAAATGATTGTCGG |
| ZCF13-AMPR-NotI | TCAGCGGCCGCTTTAGTGTCAGGTGTTAACACAACA |
| ZCF13-FLPINT-F | CAAGCCTCCTGTACCACCACCA |
| ZCF13-FLPINT-R | GCTTTAGTGTCAGGTGTTAACACAACACT |
| ZCF13-DET-F | CCACAACTGCAACAATCACAACAT |
| ZCF13-DET-R | CGATGATGGAGCTGTTTGATCAGAT |
| ZCF13-SEQ1-F | GTATCCCCCTCAACTAGCAGTTAG |
| ZCF13-SEQ2-F | CACCACCTTCTGTAACGACACCA |
| ZCF13-SEQ3-F | GCCAATTTCACTGATGCATTTGACATGA |
| ZCF13-SEQ4-F | CCGCACGTTTCAGAAGATCCC |
| ZCF13-SEQ5-F | GCCGCTAGTGATCAACTGTTTTTCC |
| Neut5homologyF | GCAGATATGAGATAAAAGTTTTAAAGGACAAGAAAAGG |
| RBK1CC9KO-F | ACAAAGTTTCTAGTATACTTAAAGTTGTGTTATTAAGTAGGGTTTTCCCAGTCACGACGT |

| | |
|---|---|
| RBK1CC9KO-R | ATAGTCAGGCAGAAGAGTTGTATTTAGAGAGGTATGACGGTGTGGAATTGTGAGCGGAT |
| RBK1DETF | GGTGATGATTCATTTGGTCAACAATTG |
| RBK1DETR | CGCCAGTAGAGTTTGTGACATTAC |
| RBK1INTF | GAGCTGAATTGGAAGCTGCTG |
| RBK1INTR | GTGCTGGTACTATGATTGCTGC |
| FLPINTF | CGCGCGTAATACGACTCACT |
| FLPINTR | CAAGCGCGCAATTAACCCTC |
| RBK1_Pr_OL-F | CCTCGAGGTCGACGGTATCGGTAGAGATGTTGAACCGAACCTAATGG |
| RBK1_ORF-OL-R | ATTTGCTTAGCATGCACGCGCTACAAAATTTCCATTACTTCTTCATAACT |
| NEUT5homology-pDISF | AGGAGGCTCCCCAAAGATTTTATCA |
| NEUT5homology-pDISR | TAACACACTGAATTCTACATCGAACAAGAGAAA |
| Nt5ADHup-UNIVOL-R | CGATACCGTCGACCTCGAGG |
| ADH1t-UNIVOL-F | CGCGTGCATGCTAAGCAAAT |
| NAT1INTF | CCCAGATGCGAAGTTAAGTGCG |
| NEUT5LAMPR | GGAATTTCTAGTCACTTGACACGACC |
| RBK1_ORF-F-SalI | TCA<u>GTCGAC</u>ATGTCATCCTCCAATCCATCAATTACTAT |
| RBK1_ORF-R-MluI | TCA<u>ACGCGT</u>CTACAAAATTTCCATTACTTCTTCATAACT |
| PrTEF1OLF | CCTCGAGGTCGACGGTATCGCTGCAAATCTGTTTGCTGATGGACA |
| tADH1OLR | ATTTGCTTAGCATGCACGCG |
| HGT7_ORF-F-SalI | TCA<u>GTCGAC</u>ATGTCTCAAGACAACGTCTCATCA |
| HGT7_ORF-R-MluI | TCA<u>ACGCGT</u>TTAAACGTGTTCCTCTTCTGGTTT |
| HGT7DETF | ACAGCTGAGGCTGTAAATAATGAAA |
| NAT1_INTR | CCATCCAAAGCTTCAATAGCTTCAGC |
| ADHterm-OL-R | CGAAAACTTGAAACTTGAAAACACCG |
| PrHygR-OL-F | CGGTGTTTTCAAGTTTCAAGTTTTCGCCATCATAAAATGTCGAGCGTCAAAAC |
| tHygR-OL-R | CTGCAGAGGACCACCTTTGATT |
| NEUT5L_OL-F | AATCAAAGGTGGTCCTCTGCAGCAGTGTCGACGTTTAAACGAGCTC |
| ADH13SEQR | ATATCGCACTCACGTAAACAC |
| NEUT5LAMPF | GCTGAATCACTTGATAGGATTTAGTTCCATTATGG |
| CaHygB-SEQR | CCAAATGTCTAACTTCTGGACAATCTTCA |
| QPCR_RBK1_F | CAAACTTCAGGAGTTGCGGTG |

| | |
|---|---|
| QPCR_RBK1_R | GGTGGTGGTGATGCTTCCAG |
| QPCR_ZCF13_F | CACCGCAAGAAAGCGAATCA |
| QPCR_ZCF13_R | GTTGCCATTCATTACCCGTTGA |
| QPCR_HGT7_F | ACAGCTGAGGCTGTAAATAATGAAA |
| QPCR_HGT7_R | ACCACCAAAGGCAATAAGGAAAC |
| QPCR_HGT8_F | AAGAGTCTTTGTGGGTGCCA |
| QPCR_HGT8_R | ACACCAATAATGATGGACGTTTGG |
| CaACT1-F-qPCR | TTGGATTCTGGTGATGGTGTTA |
| CaACT1-R-qPCR | TCAAGTCTCTACCAGCCAAATC |
| **crRNAs** | |
| crRBK1up | AGATGTTGAACCGAACCTAA |
| crRBK1down | GTTGTTGTCGCTAGTGTTTA |
| crNEUT5L-up_pDIS3 | GCTCGGAGGAGGCTCCCCAA |

**Supplementary Table 3: Oligonucleotides and crRNAs used in this work.** Point mutations are bolded and italicized. Restriction enzyme sites are underlined.

| Plasmid | Relevant characteristics | Reference |
|---|---|---|
| pDB22 | *agr* P3-GFPmut2 reporter; Erm$^{R}$ | (17) |
| pMK4 | high copy number plasmid; Amp$^{R}$, Cm$^{R}$ | (18) |
| pSK5630 | low copy number plasmid; Amp$^{R}$, Cm$^{R}$ | (19) |
| pSK-hla | pSK5630 containing Pr*hla-hla* amplified from JE2 | (2) |
| pSK-H35L | mutagenesis of pSK-hla using primers H35L-SuperFi-F and H35L-SuperFi-R | this study |
| pMV306G13+Lux | contains *luxABCDE* operon from *Mycobacterium marinum* and *Photorhabdus luminescens*; Kan$^{R}$ | (20) |
| pOLux1 | NotI and NcoI digested pMV306G13+Lux containing *agr* P3 promoter amplified from synthetic DNA *agr*P3 with primers SynthGeneF and SynthGeneRv4 | this study |
| pOLux2 | pMK4 containing *agr P3-luxABCDE* amplified from pOLux1 using primers luxABCDE-Fv2-EcoRI + luxABCDE-R-NarI | this study |
| pOLux3 | Acc65I-digested pOLux2 containing the *sarA* P1 promoter amplified from synthetic DNA *sarA*P1 using primers SynthGeneF and SynthGeneRv4 | this study |
| pBSS2 | *SAT1*-flipper plasmid; NAT$^{R}$ | (3) |
| pBSS2-ZCF13-F | plasmid pBSS2 containing 5’ flanking region of ZCF13 amplified from SC5314 gDNA using primers ZCF13-FF-KpnI and ZCF13-FR-ApaI | this study |
| pBSS2-ZCF13-FR | plasmid pBSS2-ZCF13-F 3’ flanking region of ZCF13 amplified from SC5314 gDNA using primers ZCF13-RF-NotI and ZCF13-RR-SacI | this study |
| pDUP3 | contains *NEUT5L* homology regions; NAT$^{R}$ | (21) |
| pDUP3-ZCF13 | pDUP3 containing *C. albicans ZCF13* locus between SmaI and NotI restriction sites | this study |
| CaHygB-flipper | *C. albicans* optimized HygB-flipper plasmid; HygB$^{R}$ | (7) |
| pDIS3-tADH1 | contains *NEUT5L* homology regions and *C. albicans ADH1* terminator; NAT$^{R}$ | (7) |
| pKE4 | contains *C. albicans TEF1* promoter, *ADH1* terminator, and *URA3* marker | (22) |
| pKE4-RBK1 | pKE4 containing *C. albicans* RBK1 open reading frame between SalI and MluI restriction sites | this study |
| pKE4-HGT7 | pKE4 containing *C. albicans* HGT7 open reading frame between SalI and MluI restriction sites | this study |

**Supplementary Table 4. Plasmids constructed or used in this study.** Sensitivity ($^{S}$) or resistance ($^{R}$) to the following antimicrobials is annotated: ampicillin (Amp), erythromycin (Erm), chloramphenicol (Cm), kanamycin (Kan), nourseothricin (NAT), hygromycin B (HygB).

| **Synthetic DNA** | **Sequence 5'→3'** |
|---|---|
| *sarA*P1 | CGCAGTTACGGATCAGTCACGGTACCCTGATATTTTTGACTAAACCAAATGCTAACCCAGAA<br>ATACAATCACTGTGTCTAATGAATAATTTGTTTTATAAACACTTTTTTGTTTACTTCTCATTTTT<br>AATTAGTTATAATTAACTAAATAATAGAGCATTAAATATATTTAATAAAACTTATTTAATGCAAA<br>ATTATGACTAACATATCTATAATAAATAAAGATTAGATATCAATATATTATCGGGCAAATGTAT<br>CGAGCAAGATGCATCGGTACCAGCAAAGAACATGGCAGCACC |
| *agr*P3 | CGCAGTTACGGATCAGTCACGCGGCCGCATTTTAACATAAAAAAATTTACAGTTAAGAATAA<br>AAAACGACTAGTTAAGAAAAATTGGAAAATAAATGCTTTTAGCATGTTTTAATATAACTAGATT<br>CCATGGAGCAAAGAACATGGCAGCACC |

**Supplementary Table 5. Synthetic DNA sequences used in this study.**

| Sample | Description | Total Reads Obtained | # Reads Mapped to Ref Genome | % Reads Mapped to Ref Genome |
|---|---|---|---|---|
| ***C. albicans* samples** | | | | |
| TF_WT_1 | TF library control strain (rep 1) | 40,668,310 | 39,306,969 | 96.7% |
| TF_WT_2 | TF library control strain (rep 2) | 55,404,172 | 53,547,217 | 96.7% |
| TF_WT_3 | TF library control strain (rep 3) | 39,481,300 | 38,322,397 | 97.1% |
| TF_Z_1 | TF *zcf13*Δ/Δ (rep 1) | 41,994,578 | 40,791,150 | 97.1% |
| TF_Z_2 | TF *zcf13*Δ/Δ (rep 2) | 49,034,592 | 47,232,306 | 96.3% |
| TF_Z_3 | TF *zcf13*Δ/Δ (rep 3) | 44,139,832 | 42,910,918 | 97.2% |
| ***S. aureus samples*** | | | | |
| SCTRL_1 | untreated control | 16,651,994 | 16,588,211 | 99.6% |
| SCTRL_2 | untreated control | 18,145,960 | 18,075,554 | 99.6% |
| SCTRL_3 | untreated control | 19,531,768 | 19,458,825 | 99.6% |
| SCTRL_4 | untreated control | 17,597,256 | 17,530,560 | 99.6% |
| SRIBH_1 | ribose treated | 14,238,240 | 14,180,783 | 99.6% |
| SRIBH_2 | ribose treated | 15,223,534 | 15,156,847 | 99.5% |
| SRIBH_3 | ribose treated | 16,051,650 | 15,985,920 | 99.6% |
| SRIBH_4 | ribose treated | 13,630,960 | 13,579,086 | 99.6% |

**Supplementary Table 6. RNA-seq mapping statistics in each sample and sample key.**

**SUPPLEMENTARY FIGURES**

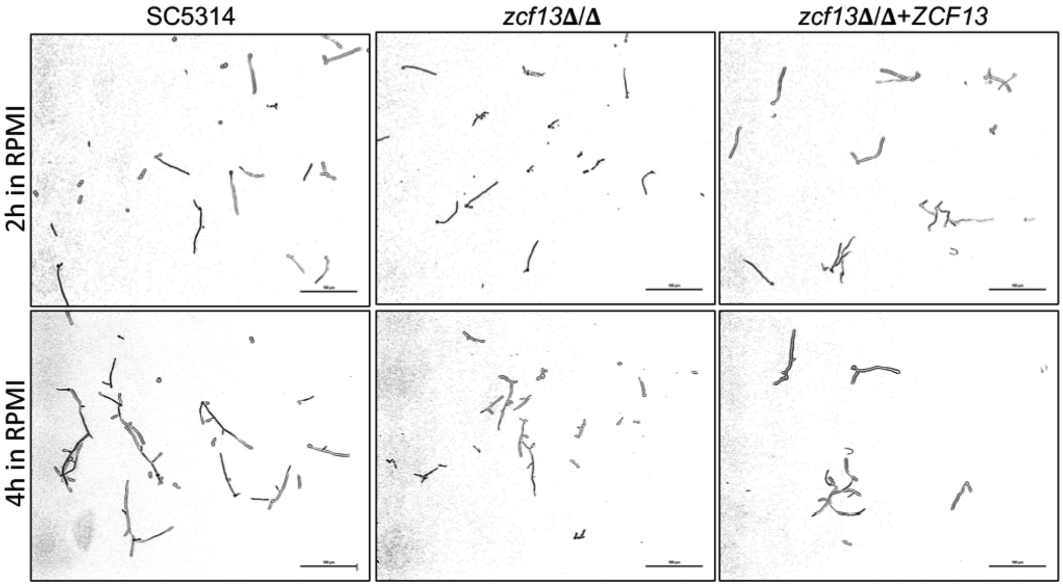


**Supplementary Fig 1:** *ZCF13* deletion does not impact hyphal growth. SC5314, *zcf13***Δ**/**Δ**, and *zcf13***Δ**/**Δ**+*ZCF13* cells were inoculated in RPMI containing 165 mM MOPS, pH 7.0 at 37ºC, and hyphal formation was monitored at 2 h and 4 h. Images were digitally captured by standard light microscopy. Scale bar is 100 µm. Images are representative of duplicate experiments.

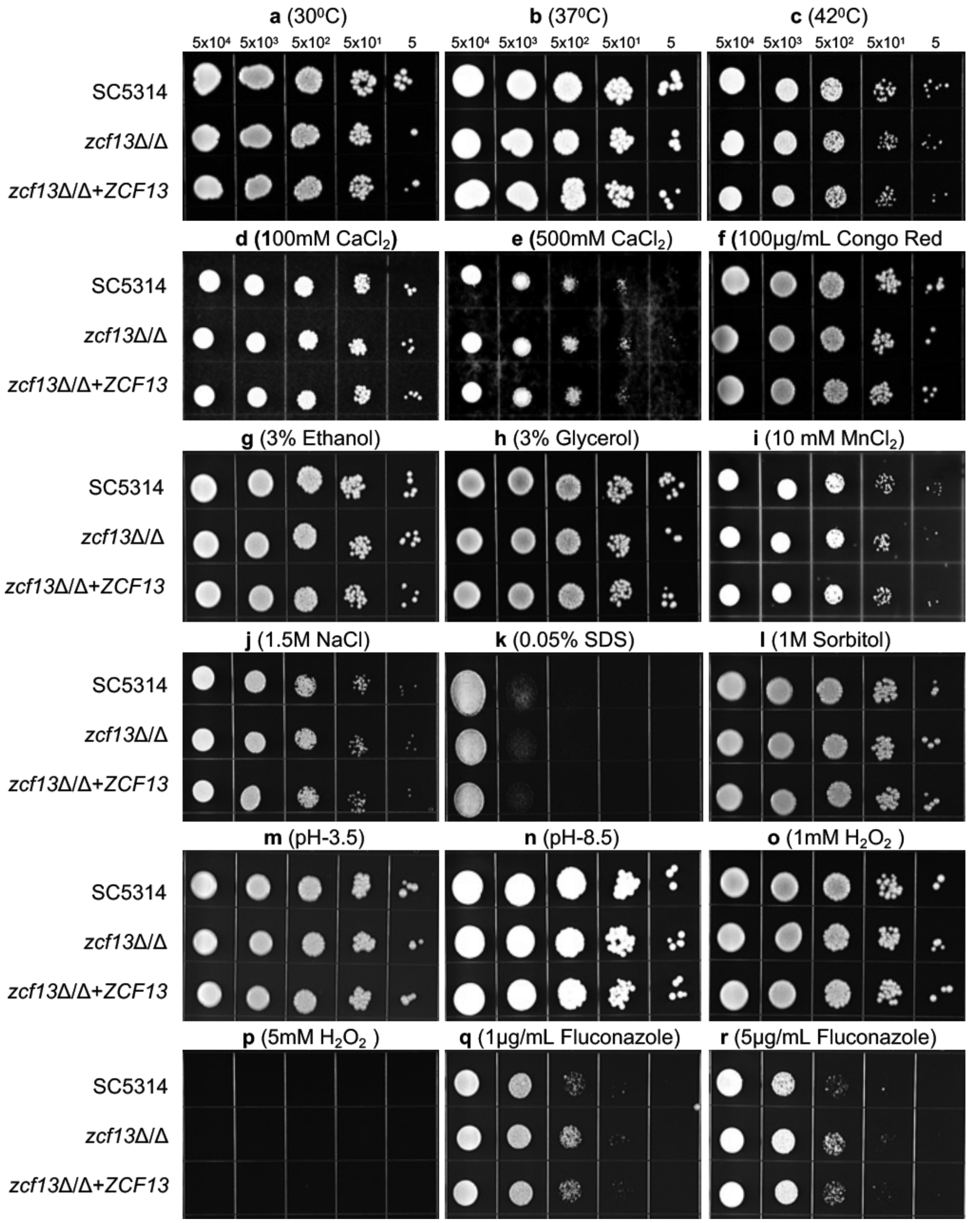


**Supplementary Fig. 2: *ZCF13* deletion does not increase susceptibility to a variety of stressors.** The indicated number of SC5314, *zcf13*Δ/Δ, *zcf13*Δ/Δ+*ZCF13* cells were spotted on YPD medium containing various stressors or incubated under varied temperature as indicated: **a**, **b, c** temperature, **d, e** CaCl2, **f** Congo Red, **g** ethanol, **h** glycerol, **i** MnCl2, **j** NaCl, **k** SDS, **l** sorbitol, **m, n** pH, **o, p** H2O2, **q, r** fluconazole. Images were digitally captured. Images are representative of duplicate experiments.

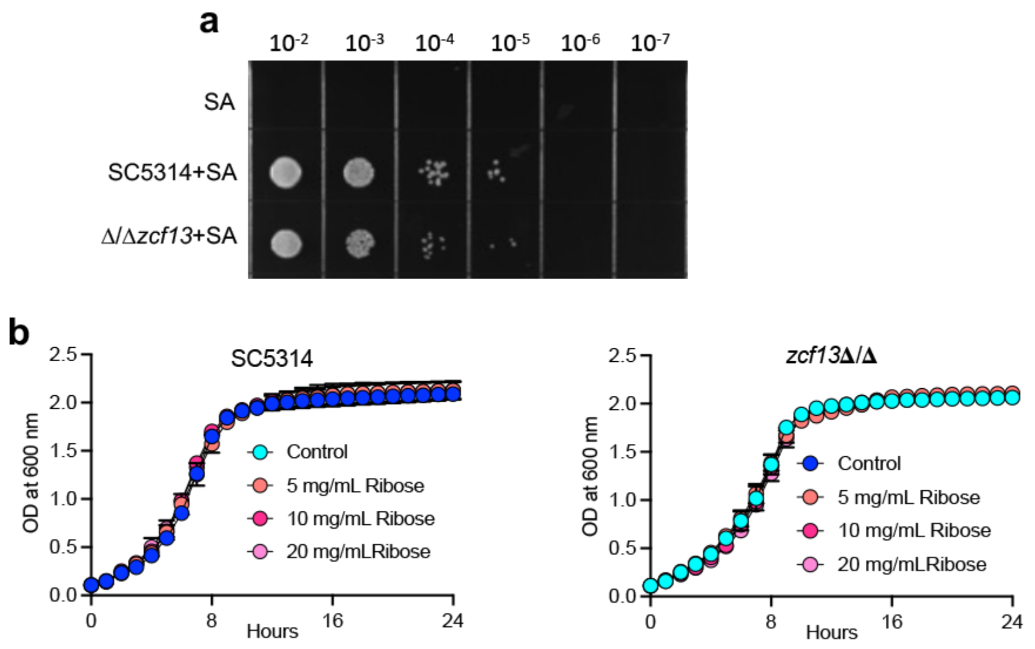


**Supplementary Fig. 3: Ribose is not growth inhibitory for *C. albicans*. a**. SC5314, *zcf13*Δ/Δ and *zcf13*Δ/Δ+ZCF13 were cocultured with S. aureus(pDB22) in TSBg supplemented with 0.5 mg/mL ribose for 16 h. Indicated dilutions were made, spotted onto YPD containing 50 µg/mL chloramphenicol, and cell growth monitored at 24 h. Images are representative of 3 independent experiments. **b** SC5314 and zcf13Δ/Δ were grown in TSBg supplemented with indicated concentrations of ribose for 16 h and growth temporally monitored at OD600 nm using a spectrophotometer. Data is depicted as the mean ± SEM of 3 independent experiments.

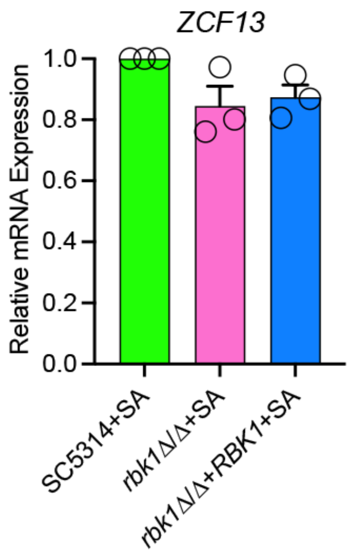


**Supplementary Fig. 4: *RBK1* deletion does not impact *ZCF13* expression.** SC5314 (green), *rbk1*Δ/Δ (pink), *rbk1*Δ/Δ and *rbk1*Δ/Δ+*RBK1* (blue) were co-cultured with *S. aureus*(pDB22) in TSBg. *ZCF13* expression was measured by quantitative real-time PCR and normalized by $2^{-\Delta\Delta Ct}$ to *ACT1* and SC5314+SA using gene-specific primers**.** Data is depicted as the mean + SEM from 3 independent experiments.

```
name                          sequence                               position

ScGal2p       MAVEENNMPVVSQQPQAGEDVISSLSKDSHLSAQSQKYSN--DELKAGESGSEGSQSVPI      58
CaHgt7p       ----------MSQ------------DNVSSTSTA----EAVNNEIKV-KDEFPQEEQAHT      33
CaHgt8p       ----------MSS------------TNSTENHAVEEKYEDPQQQQQQ-QQQQQQQQQQKD      37
                        :*.             : :   :     .   :: :  :.    .:.

ScGal2p       EIPKKPMSEYVTVSLLCLCVAFGGFMFGWDTGTISGFVVQTDFLRRFGMKHKDGTHYLSN      118
CaHgt7p       SLEDKPVSAYIGIIIMCFLIAFGGFVFGFDTGTISGFINMSDFLERFGGTKADGTLYFSN      93
CaHgt8p       ALAKKPMSAYIGISIMCVLIAFGGFVFGFDTGTISGFINMSDFLERFGGTRADGTLYFSN      97
               : .**:* *: : ::*. :*****:**:********:  :***.*** .: *** *:**

ScGal2p       VRTGLIVAIFNIGCAFGGIILSKGGDMYGRKKGLSIVVSVYIVGIIIQIASINKWYQYFI      178
CaHgt7p       VRTGLMIGLFNAGCAIGALFLSKVGDMYGRRVGIMTAMIVYIVGIIVQIASQHAWYQVMI      153
CaHgt8p       VRTGLLIGLFNVGCAIGALFLSKVGDMYGRRVGIMTAMIIYIVGIIVQIASQHAWYQVMI      157
              *****::.:** ***:*.::*** ******: *:  .: :******:**** : *** :*

ScGal2p       GRIISGLGVGGIAVLCPMLISEIAPKHLRGTLVSCYQLMITAGIFLGYCTNYGTKSYSNS      238
CaHgt7p       GRIITGLAVGMLSVLCPLFISEVSPKHLRGTLVCCFQLMITLGIFLGYCTTYGTKSYSDS      213
CaHgt8p       GRIITGLAVGTLSVLCPLFISEVSPKHLRGTLVCCFQLMITLGIFLGYCTTYGTKTYSDS      217
              ****:**.** ::****::***::*********.*:***** ********.****:**:*

ScGal2p       VQWRVPLGLCFAWSLFMIGALTLVPESPRYLCEVNKVEDAKRSIAKSNKVSPEDPAVQAE      298
CaHgt7p       RQWRIPLGLCFAWALCLVAGMVRMPESPRYLVGKDRIEDAKMSLAKTNKVSPEDPALYRE      273
CaHgt8p       RQWRIPLGLCFAWALCLLGGMVRMPESPRYLVGNDRIEDAKISLAKTNKVSPEDPALYRE      277
               ***:********:* ::..:. :*******   :::**** *:**:*********:  *

ScGal2p       LDLIMAGIEAEKLAGNASWGELFSTKTKVFQRLLMGVFVQMFQQLTGNNYFFYYGTVIFK      358
CaHgt7p       LQLIQAGVERERLAGKASWGTLFNGKPRIFERVIVGVMLQALQQLTGDNYFFYYSTTIFK      333
CaHgt8p       LQLIQAGVERERLAGSASWKSLITGKPRILERVFVGAMLQSLQQLTGDNYFFYYSTTIFK      337
              *:** **:* *:***.***  *:. * ::::*::*.::* :*****:******.*.***

ScGal2p       SVGLDDSFETSIVIGVVNFASTFFSLWTVENLGHRKCLLLGAATMMACMVIYASVGVTRL      418
CaHgt7p       SVGMNDSFETSIIIGVINFASTFVGIYAIERMGRRLCLLTGSVAMSICFLIYSLVGTQHL      393
CaHgt8p       SVGLNDSFQTSIIIGVINFASTFVGIYAIEKLGRRHCLLIGSVAMSICFLIYSLIGTQHL      397
              ***::***:***:***:******..::::*.:*:* *** *:.:*  *::**: :*. :*

ScGal2p       YPHGKSQPSSKGAGNCMIVFTCFYIFCYATTWAPVAWVITAESFPLRVKSKCMALASASN      478
CaHgt7p       YIDKPGGASRKPDGDAMIFMTSLYVFFFASTWAGGVYSIISELYPLKVRSKAMGLANASN      453
CaHgt8p       YIDKPGGASRKPDGDAMIFITALYVFFFASTWAGGVYSIMSELYPLKVRSKAMGIASASN      457
              * .  .  * *  *:.**.:*.:*:* :*:***  .: * :* :**:*:**.*.:*.***

ScGal2p       WVWGFLIAFFTPFITSAINFYYGYVFMGCLVAMFFYVFFFVPETKGLSLEEIQELWEEGV      538
CaHgt7p       WTWGFLISFFTSFITDAIHFYYGFVFMGCLVFSIFFVYFMVYETKGLTLEEIDELYSTKV      513
CaHgt8p       WLWGFLISFFTSFITDAIHFYYGFVFMGCLVFSIFFVYFMVYETKGLTLEEIDELYSTKV      517
              * *****:*** ***.**:****:*******  :*:*:*:* *****:****:**:.  *

ScGal2p       LPWKSEGWIPSSRRGNNYDLEDLQHDDKPWYKAMLE*      574
CaHgt7p       LPWKSAGWVPPSEEEMATSTG-YAGDAKPEEEHV*--      546
CaHgt8p       VPWKSAGWVPPSEEEMATSTG-YAGDAKPTEEHV*--      550
              :**** **:* *..    .      * **  : :
```

**Supplementary Fig. 5: *C. albicans* Hgt7p and Hgt8p are close orthologues to the *S. cerevisiae* low affinity ribose importer Gal2p.** A multiple sequence alignment was performed using Clustal Omega. “*” indicate exact matches; “:” or “.” indicate matches to amino acids with strongly or weakly similar properties, respectively.
